# Neurohormonal signalling controls insulin sensitivity and specificity in *C. elegans*

**DOI:** 10.1101/229393

**Authors:** Nicholas O. Burton, Vivek K. Dwivedi, Kirk B. Burkhart, E.W. Rebecca Kaplan, L. Ryan Baugh, H. Robert Horvitz

## Abstract

Insulin and insulin-like growth factor signalling regulates a broad spectrum of growth and metabolic responses to a variety of internal and environmental stimuli. Such responses can be tailored to the environment so that changes in insulin signalling result in distinct physiological responses to different stimuli. For example, the inhibition of insulin-like signalling by osmotic stress or by starvation of *C. elegans* results in physiologically distinct states and patterns of gene expression. How does insulin-like signalling elicit different responses to different environmental stimuli? We report that neurohormonal signalling involving the *C. elegans* cytosolic sulfotransferase SSU-1 controls developmental arrest in response to osmotic stress but not to starvation; that SSU-1 functions in a single pair of sensory neurons to control signalling via the nuclear hormone receptor NHR-1; that signalling controlled by SSU-1 acts antagonistically to insulin-like signalling; and that the FOXO transcription factor DAF-16, a downstream effector of insulin-like signalling, enters the nucleus in response to osmotic stress but activates gene expression only if SSU-1 is active. We propose that neurohormonal signalling controlled by one or more cytosolic sulfotransferases similarly regulates the specificity of responses to changes in insulin signalling during periods of environmental stress in other organisms and that abnormalities in such sulfotransferase-controlled neurohormonal signalling might contribute to human disorders that involve insulin signalling, such as obesity and type 2 diabetes.

---

We previously reported that *C. elegans* arrests its development immediately after hatching in response to osmotic stress and that this developmental arrest is caused by the inhibition of insulin-like signalling and subsequent activation of the FOXO transcription factor DAF-16^1^. Other studies showed that *C. elegans* also arrests its development immediately after hatching in response to starvation and that this developmental arrest is similarly caused by the inhibition of insulin-like signalling and activation of DAF-16^2^. Despite these similarities in the timing and regulation of developmental arrest in response to osmotic stress and starvation, we found that these two developmental arrests are physiologically distinct^1^. Specifically, (a) developmental arrest in response to osmotic stress results in animals that are immobile and do not respond to touch, whereas animals arrested in response to starvation remain mobile, and (b) most genes that exhibit changes in expression in animals arrested in response to osmotic stress do not exhibit changes in expression in animals arrested in response to starvation^1^. These results suggest that the inhibition of insulin-like signalling can elicit different physiological responses to starvation and osmotic stress. The mechanistic basis of these different responses is unknown.

To determine how the inhibition of insulin-like signalling can result in distinct states of arrested development in response to osmotic stress and starvation, we screened for mutants that failed to arrest development in response to osmotic stress (500 mM NaCl). We identified a nonsense allele of the cytosolic sulfotransferase *ssu-1* (*n5888* W284Stop) that resulted in approximately 40% of animals failing to arrest development in response to 500 mM NaCl (Fig. 1a and 1b). Similarly, six independently isolated mutations in *ssu-1* (*fc73, tm1117, gk266317, gk747222, gk876992, gk319712*) caused animals to fail to arrest development in response to osmotic stress (Fig. 1b). SSU-1 is expressed in a single pair of sensory neurons, the ASJ neurons^3^. To determine if SSU-1 functions specifically in the ASJ sensory neurons to regulate developmental arrest in response to osmotic stress we expressed a rescuing transgene *of ssu-1(+)* under the control of the ASJ-specific promoter *trx-1* in *ssu-1(-)* animals^6^. Expression of SSU-1 in the ASJ sensory neurons restored developmental arrest in response to osmotic stress (Fig. 1c), indicating that SSU-1 functions in the ASJ sensory neurons. We conclude that SSU-1 promotes developmental arrest in response to osmotic stress. SSU-1 is the only predicted cytosolic sulfotransferase encoded in the *C. elegans* genome^3^ and is most similar to the human cytosolic sulfotransferase SULT2A1, which sulfonates a variety of endogenous steroids, including dehydroepiandrosterone^4^. We propose that SSU-1 controls neurohormonal signalling by sulfonating signalling molecules in the ASJ sensory neurons to regulate developmental arrest in response to osmotic stress.

**Fig. 1.**
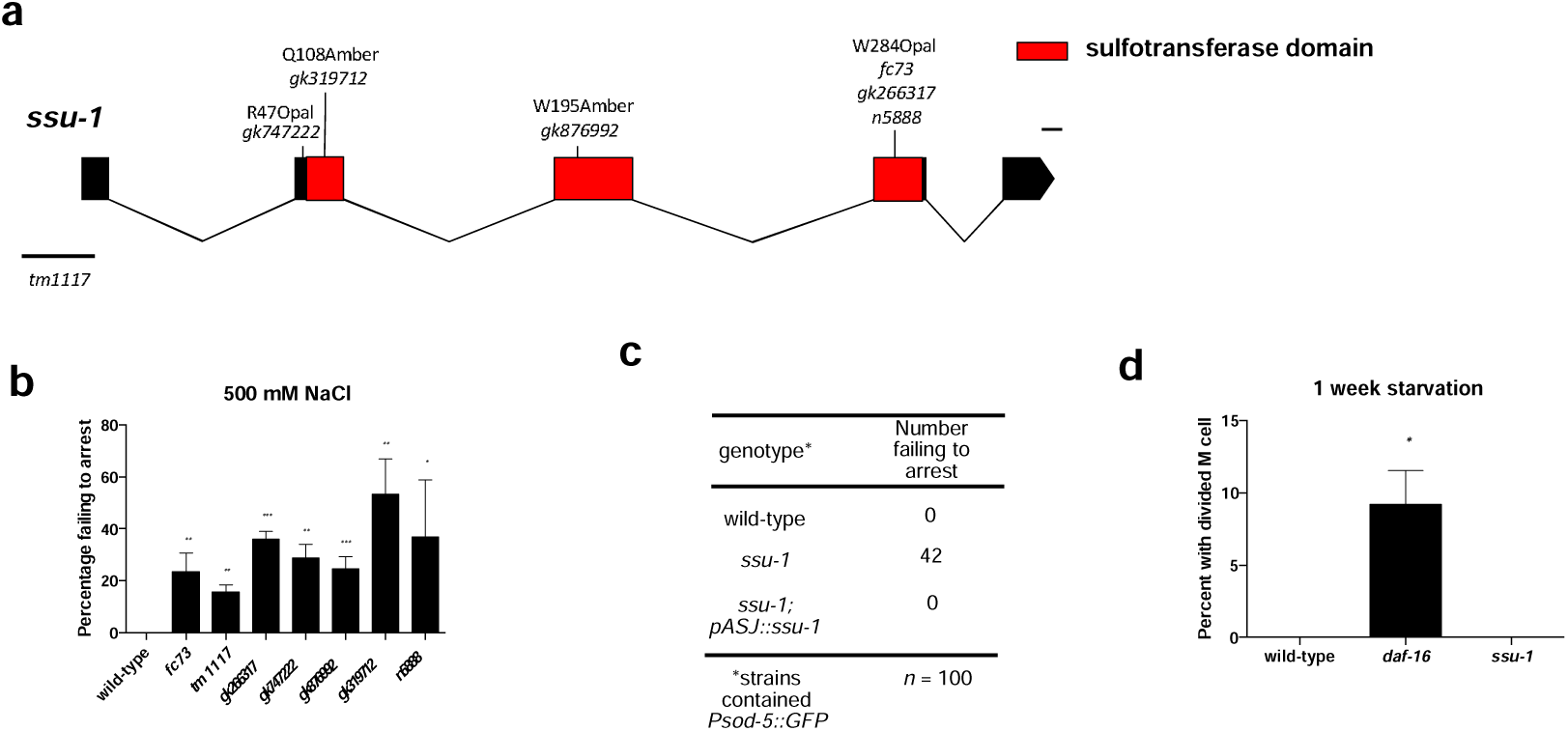
The cytosolic sulfotransferase SSU-1 functions in the ASJ sensory neurons to regulate developmental arrest in response to osmotic stress, (a) Schematic of *ssu-1* mutations that cause defects in developmental arrest in response to osmotic stress. Scale bar 100 bp (b) Percent of *ssu-1* mutants failing to arrest development in response to 500 mM NaCl. n = 3 experiments represent greater than 100 animals. Error bars, s.d. (c) Number of wild-type, *ssu-1 (fc73)*, and *ssu-1 (fc73)*; *ssu-1 (+)* animals failing to arrest development in response to osmotic stress. The *trx-1* promoter was used to drive ASJ cell specific expression of *ssu-1.* Animals also contained *wuIs57 [pPD95.77 Psod-5::GFP, rol-6(su1006)]* which drives GFP expression in response to osmotic stress, n = 100 animals (d) Percent of wild-type, *daf-16(mu86)*, and *ssu-1 (fc73)* animals with a divided M cell after 1 week of starvation, n > 200 animals. Error bars, s.e.m. * p < 0.05, ** p < 0.01, *** p < 0.001.

To test if SSU-1 is also required for developmental arrest in response to starvation, we starved wild-type, *daf-16* and *ssu-1* mutant animals for one week and assayed the percentage of animals with a divided M cell; M cell division does not occur in animals that arrest development in response to starvation^2^. We found that 100% of M cells in both wild-type animals and *ssu-1* mutants arrested cell division (Fig. 1d). By contrast, 8% of M cells in *daf-16* mutants failed to arrest cell division in response to starvation (Fig. 1d), consistent with previous observations^2^. These results indicate that SSU-1 is required for developmental arrest in response to osmotic stress but is not required for developmental arrest in response to starvation.

In humans, the cytosolic sulfotransferase SULT2A1 sulfonates steroid hormones such as dehydroepiandrosterone (DHEA) and pregnenolone^4^. These hormones in turn regulate gene expression by activating nuclear hormone receptors^4^. We hypothesized that SSU-1 might similarly regulate the sulfonation of a hormone that in turn promotes developmental arrest in response to osmotic stress by controlling the transcriptional response to osmotic stress. To test if SSU-1 regulates gene expression in response to osmotic stress, we exposed wild-type and *ssu-1* mutant embryos to either 50 mM or 500 mM NaCl for 3 hours and quantified mRNA expression using RNA-Seq. We found that the expression of 434 genes was upregulated greater than 2-fold in response to osmotic stress and that SSU-1 function was required for the expression of 106 of these genes (Supplemental Table 1), including 20 of the 25 genes that exhibited a greater than 10-fold increase in expression in response to osmotic stress (*p* < 0.05) (Fig. 2a). For example, the genes that encode the superoxide dismutase *sod-5* and the osmotic stress resistance protein *lea-1*^6^ exhibited a greater than 25-fold increase in expression in response to osmotic stress and their increased expression in response to osmotic stress required SSU-1 (Fig. 2a). We confirmed that SSU-1 was required for the increased expression of *sod-5* in response to osmotic stress using a GFP reporter (Figs. 2b and 2c). GFP was expressed broadly throughout the animal in response to osmotic stress, and this broad increase in expression required SSU-1 function in the ASJ sensory neurons (Figs. 2b and 2c).

**Fig. 2.**
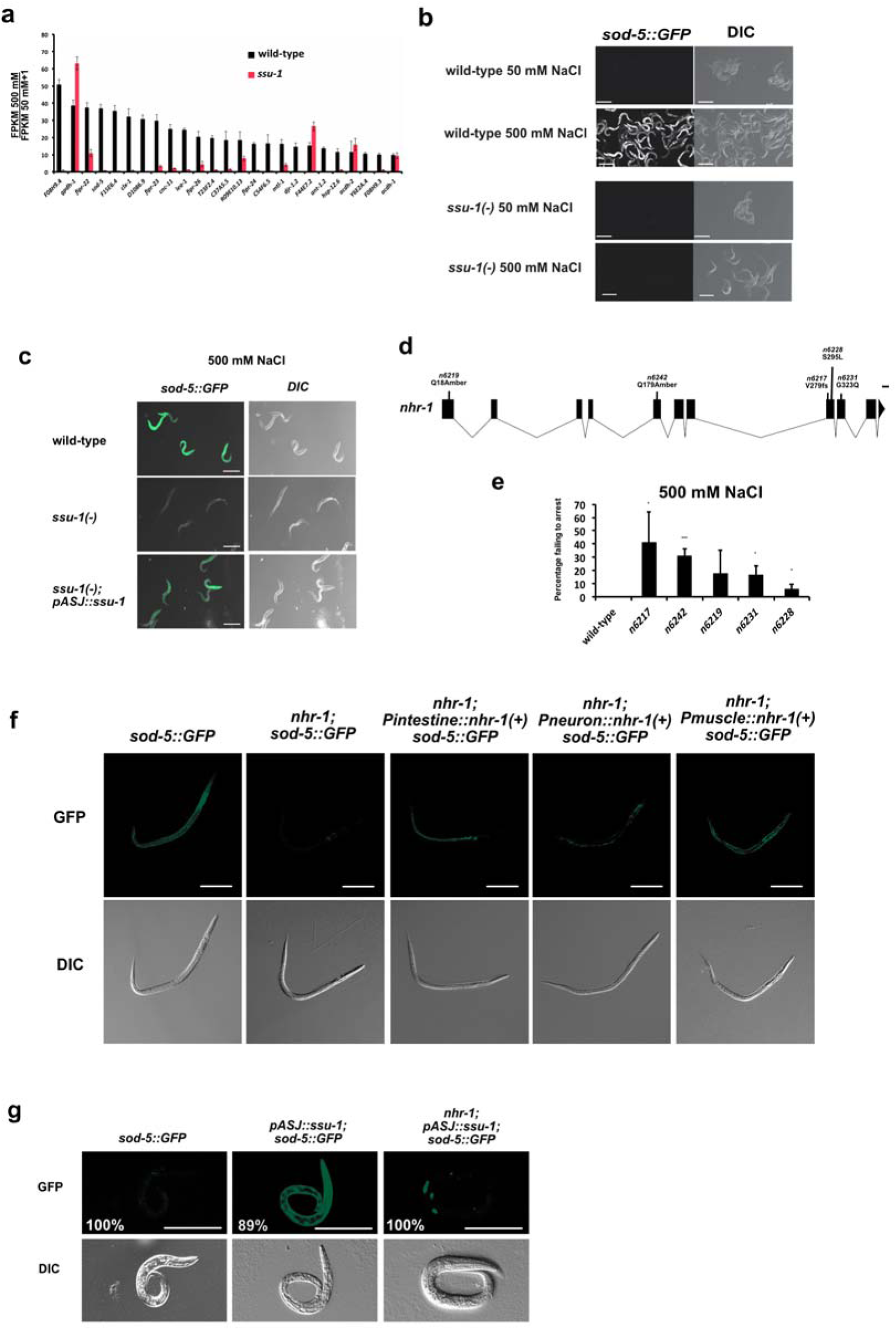
The nuclear hormone receptor NHR-1 is required for SSU-1 to promote the transcriptional response to osmotic stress (a) Average fold change mRNA expression in wild-type and *ssu-1 (fc73)* mutant animals at 500 mM NaCl when compared to 50 mM NaCl measured by RNA-Seq (FPKM 500 mM NaCl/FPKM 50 mM NaCl + 1). Shown are the 25 genes that exhibited the greatest increase in expression in wild-type animals at 500 mM NaCl vs. 50 mM NaCl. n = 3. Error bars s.d. (b) Confocal and differential interference contrast (DIC) images showing *Psod-5::gfp* expression in wild-type and *ssu-1 (fc73)* mutants exposed to 500 mM NaCl for 24 hrs. Scale bars, 100 |im. (c) Confocal and DIC images of *Psod-5::GFP* expression in wild-type, *ssu-1 (fc73)*, and *ssu-1 (fc73)*; *ssu-1 (+)* mutants exposed to 500 mM NaCl for 24 hrs. The *trx-1* promoter was used to drive ASJ cell-specific expression of SSU-1. Scale bars, 100 μm. (d) Schematic of *nhr-1* mutations that cause defects in developmental arrest in response to osmotic stress. Scale bar 100 bp (e) Percent of *nhr-1* mutants failing to arrest development in response to 500 mM NaCl. n = 3 experiments represent greater than 100 animals. Error bars, s.d. (f) Representative confocal and DIC images of *Psod-5::GFP* expression in wild-type and *nhr-1(n6242)* mutant dauers. The *ges-1* promoter was used to drive intestine specific expression, the *rab-3* promoter was used to drive neuron specific expression, and the *myo-3* promoter was used to drive muscle specific expression. Scale bar 100 μm. (f) Representative confocal and DIC images of *Psod-5::GFP* expression in wild-type and *nhr-1 (n6242)* LI stage animals. The *trx-1* promoter was used to drive ASJ cell-specific expression of SSU-1. Animals expressed GFP specifically in coelomocytes under the control of the *unc-122* promoter as a coinjection marker. Percentages reflect the percentage of animals that express GFP similar to the representative images. Scale bar 100 μm < 0.05, ** p < 0.01, *** p < 0.001.

To identify the transcription factor that functions downstream of SSU-1 to control the transcriptional response to osmotic stress we screened for additional mutants that 1) failed to arrest development in response to osmotic stress and 2) failed to express *sod-5::gfp* in response to osmotic stress. We identified five alleles of the nuclear hormone receptor *nhr-1* (*n6217, n6219, n6228, n6231, n6242*) (Figs. 2d and 2e). In addition to failing to express *sod-5::GFP* in embryos arrested in response to osmotic stress, we found that *nhr-1* mutants failed to express *sod-5::GFP* in dauer arrested animals (Fig. 2f). The loss of *sod-5::gfp* expression in dauer animals was tissue specifically rescued in animals that expressed a wild-type copy of *nhr-1* in the intestine, using the *ges-1* promoter, neurons, using the *rab-3* promoter, and muscle, using the *myo-3* promoter (Fig. 2f). However, we were unable to restore *sod-5::gfp* expression in *nhr-1* mutant embryos using these three promoters or restore *sod-5::gfp* expression using the endogenous *nhr-1* promoter. We suspect that overexpression of *nhr-1* in embryos might promote developmental arrest and prevent the recovery of transgenic animals. We conclude that NHR-1 functions cell autonomously to promote the expression of stress response genes, such as *sod-5*, in arrested animals.

We found that overexpression of *ssu-1* in the ASJ sensory neurons was sufficient to drive the expression *of sod-5::gfp* even in the absence of osmotic stress (Fig. 2g). To test if the expression of *sod-5::gfp* in animals that overexpress *ssu-1* requires NHR-1 we examined *nhr-1* mutants that overexpressed *ssu-1*. We found that the overexpression of *ssu-1* did not result in *sod-5::gfp* expression in *nhr-1* mutants (Fig. 2g). These results indicate that NHR-1 is required for SSU-1 to drive the expression of *sod-5* and suggest that SSU-1 and NHR-1 function in the same pathway. We propose that SSU-1 sulfonates the ligand for NHR-1 in the ASJ sensory neurons and that this ligand then signals to diverse tissues to activate NHR-1 and drive the transcriptional response to osmotic stress and developmental arrest.

Osmotic stress causes developmental arrest of *C. elegans* by inhibiting insulin-like signalling, which results in the activation of the FOXO transcription factor DAF-16^1^. To determine if DAF-16 and SSU-1 regulate the expression of the same or different downstream target genes we exposed wild-type and *daf-16* mutant embryos to 50 mM or 500 mM NaCl and quantified mRNA expression by RNA-seq. We found that 161 genes both exhibited a greater than 2-fold increase in expression in response to osmotic stress and were dependent on DAF-16 for their induction (Fig. 3a-b and Supplemental Table 2). Notably, all of the top 25 genes that exhibit the largest DAF-16 dependent increase in expression in response to osmotic stress also require SSU-1 for their expression, and all of the top 25 genes that exhibit the largest SSU-1 dependent increase in expression in response to osmotic stress also require DAF-16 for their expression (Supplemental Table 2). In total, of the 161 genes regulated by DAF-16 in response to osmotic stress, 64 were also regulated by SSU-1 (Fig. 3c). By contrast, genes such as *gpdh-1*, which exhibits an approximately 50-fold increase in response to osmotic stress, did not require either SSU-1 or DAF-16 for their expression (Fig. 2a and 3a). These results indicate that most of the genes that exhibit the largest increase in expression in response to osmotic stress require both SSU-1 and DAF-16 for their expression.

**Fig. 3.**
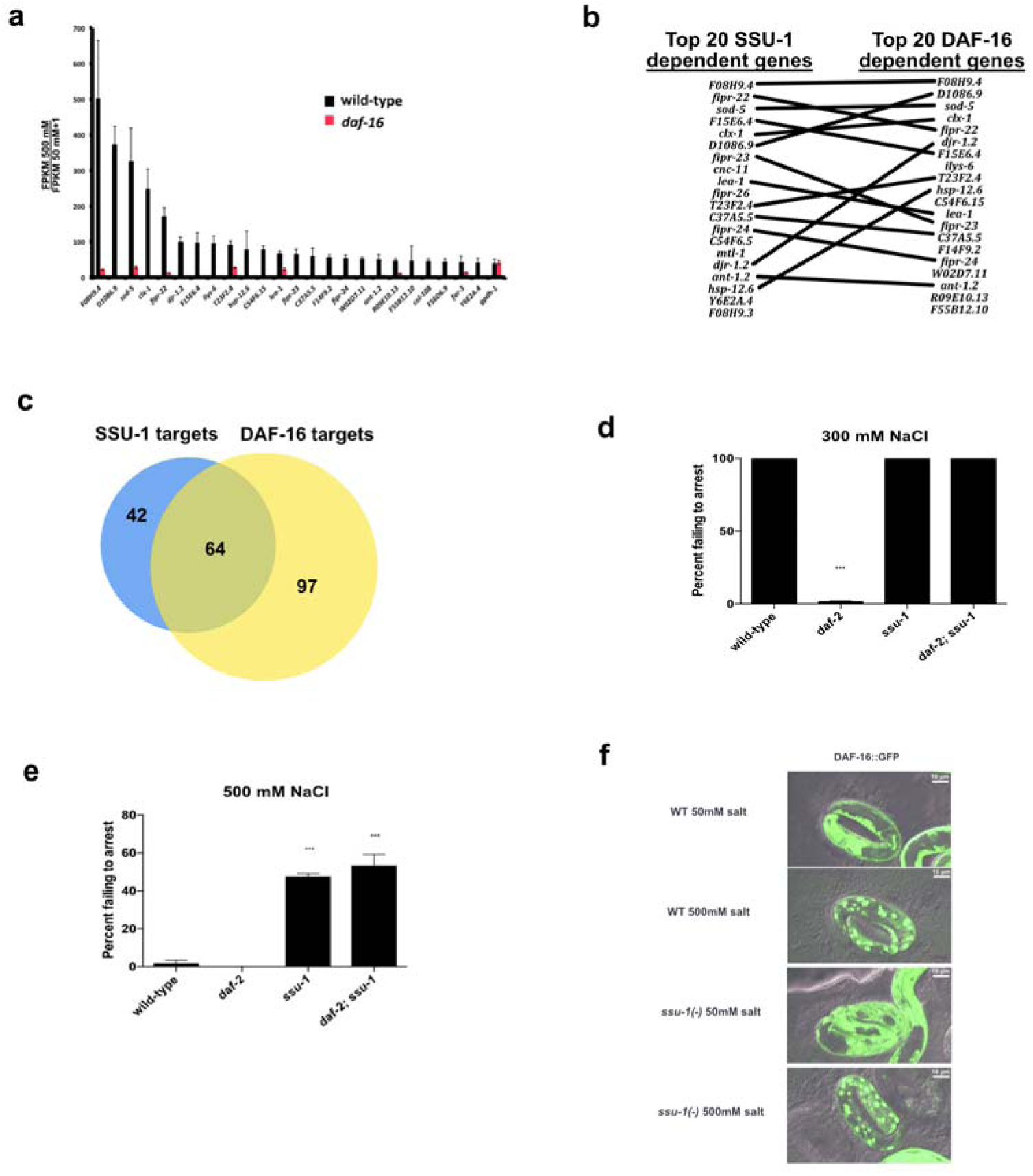
SSU-1 and insulin-like signalling via the FOXO transcription factor DAF-16 function in parallel to regulate gene expression and developmental arrest in response to osmotic stress, (a) Average fold change mRNA expression in wild-type and *daf-16(m26)* mutant animals at 500 mM NaCl when compared to 50 mM NaCl measured by RNA-Seq (FPKM 500 mM NaCl/FPKM 50 mM NaCl + 1). Shown are the 25 genes that exhibited the greatest increase in expression in wild-type animals at 500 mM NaCl vs. 50 mM NaCl. n = 3. Error bars s.d. (b) Comparison of the 20 genes exhibiting the largest SSU-1 and DAF-16 dependent increases in expression in response to 500 mM NaCl. (c) Venn diagram of genes with a greater than 2-fold increase in RNA expression in response to osmotic stress and dependent on SSU-1 (blue) and/or DAF-16 (yellow) (d) Percent of wild-type, *daf-2(e1370)* and *ssu-1(fc73)* mutants failing to arrest development in response to 300 mM NaCl. n >100 animals, error bars s.d. (e) Percent of wild-type, *daf-2(e1370)* and *ssu-1(fc73)* mutants failing to arrest development in response to 500 mM NaCl. n >100 animals. Error bars, s.d. (f) Representative confocal images of DAF-16::GFP localization after 5 hrs of exposure to osmotic stress (500 mM NaCl) in the wildtype and *ssu-1(fc73)* mutants. Scale bars, 10 μm. *** p < 0.001.

Mutations in the insulin-like receptor gene *daf-2* result in animals that are more likely to arrest development in response to osmotic stress than wild-type animals^1^. To examine potential interactions between *ssu-1* and *daf-2* we constructed double mutant animals harboring mutations in both *daf-2* and *ssu-1* and exposed wild-type, *daf-2*, *ssu-1*, and *daf-2*; *ssu-1* double mutant embryos to mild osmotic stress (300 mM NaCl) and strong osmotic stress (500 mM NaCl). Consistent with our previous findings, we found that nearly 100% of *daf-2* mutant embryos arrested development at 300 mM NaCl (Fig. 3d). However, none of the wild-type, *ssu-1*, or *daf-2*; *ssu-1* double mutant animals arrested development at 300 mM NaCl (Fig. 3d). In addition, we found that approximately 30% of *ssu-1* and *daf-2*; *ssu-1* double mutant animals failed to arrest development even at 500 mM NaCl (Fig. 3e). These results indicate that the loss of SSU-1 function can suppress the effects of reduced insulin-like signalling in response to osmotic stress.

Insulin-like signalling inhibits the activation of the FOXO transcription factor DAF-16 by sequestering it in the cytoplasm^7^. The loss of insulin-like signalling in response to osmotic stress causes developmental arrest because DAF-16 is no longer sequestered in the cytoplasm and can translocate into the nucleus^1^. We hypothesized that SSU-1 signalling might promote the translocation of DAF-16 into the nucleus in response to osmotic stress in parallel to insulin-like signalling. To test this hypothesis, we examined wild-type and *ssu-1* mutant animals that expressed a GFP-tagged copy of DAF-16 and assayed DAF-16 translocation to the nucleus in response to osmotic stress. We found that SSU-1 was not required for DAF-16 translocation into the nucleus in response to osmotic stress (Fig. 3f and Supplemental Fig. 1). This observation suggests that signalling via SSU-1 functions in parallel to insulin-like signalling and DAF-16 translocation into the nucleus to regulate development and gene expression in response to osmotic stress.

We previously found that increases in the levels of glycerol, an osmolyte that protects animals from the effects of osmotic stress, can protect *C. elegans* from developmental arrest in response to osmotic stress^1^. We hypothesized that the loss of SSU-1 or DAF-16 might similarly result in increased production of glycerol to protect animals from entering developmental arrest in response to osmotic stress. To test this hypothesis we profiled 137 polar metabolites and 1,069 lipid metabolites in wild-type, *ssu-1*, and *daf-16* mutant embryos by mass spectrometry (Supplemental Table 3). Four polar molecules and four lipid molecules were changed in both *ssu-1* and *daf-16* mutant embryos (Figs. 4a and 4b). We did not observe an increase in the levels of glycerol, indicating that the loss of SSU-1 and DAF-16 does not protect animals from developmental arrest in response to osmotic stress by increasing the production of glycerol. In addition, we found that none of the metabolites we profiled exhibited a greater than 2-fold change in abundance, suggesting that under normal osmotic conditions the loss of either SSU-1 or DAF-16 has minimal effects on metabolism.

**Fig. 4.**
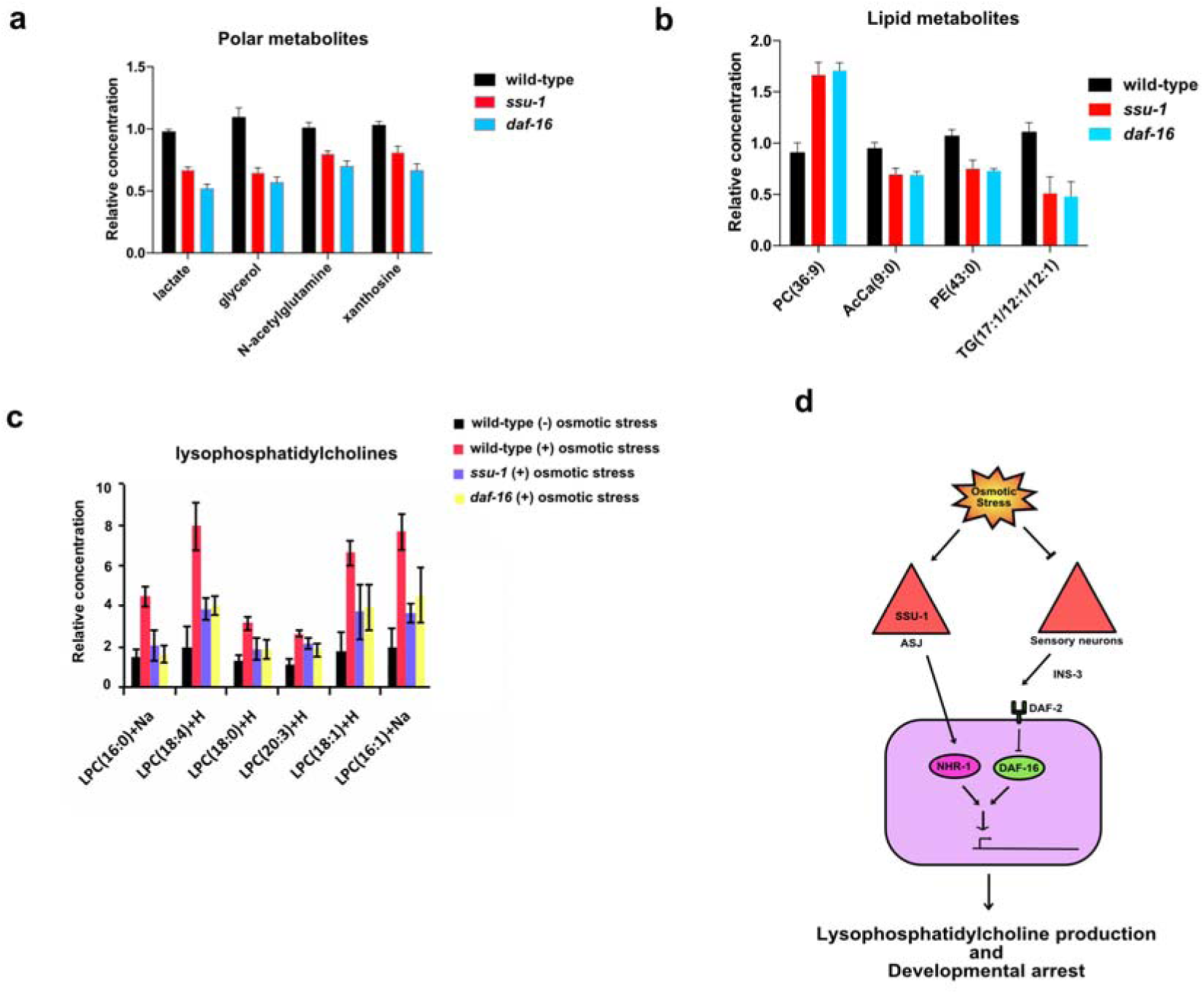
SSU-1 and DAF-16 regulate lysophosphatidylcholine levels in response to osmotic stress (a) Relative levels of polar metabolites exhibiting a statistically significant (p < 0.01) change in levels in *ssu-1(fc73)* and *daf-16(m26)* mutant embryos. Metabolites were normalized to the levels of histidine, n = 3. Error bars, s.d. (c) Relative levels of lipid metabolites exhibiting a statistically significant (p < 0.01) change in levels in both *ssu-1(fc73)* and *daf-16(m26)* mutant embryos. Metabolites were normalized to the levels of total lipid, n = 3. Error bars, s.d. PC phosphatidylcholine; AcCa Acylcarnitine; PE phosphatidylethanolamine; TG triglyceride, (c) Relative levels of lysophosphatidylcholine (LPC) metabolites in wild-type, *ssu-1(fc73)* and *daf-16(m26)* mutant embryos. Metabolites shown exhibited a statistically significant (p < 0.01) and greater than 2-fold increase in abundance in response to osmotic stress (300 mM NaCl) and required both SSU-1 and DAF-16. Metabolites were normalized to the levels of total lipid. n = 3. Error bars, s.d (d) Model for how *C. elegans* regulates metabolism and development in response to osmotic stress. See text for details.

Alternatively, SSU-1 and DAF-16 might regulate the production of glycerol specifically in response to osmotic stress, such that in response to osmotic stress *ssu-1* and *daf-16* mutants produce more glycerol, which protects animals from developmental arrest. To determine how SSU-1 and DAF-16 regulate metabolism in response to osmotic stress we profiled polar and lipid metabolites in embryos exposed to 300 mM NaCl. We identified 6 polar metabolites that increased in abundance more than 2-fold (p < 0.01) in wild-type animals in response to osmotic stress, including glycerol, and 12 polar metabolites that decreased in abundance more than 2-fold (p < 0.01) in wild-type animals in response to osmotic stress, including several TCA cycle intermediates (Supplemental Table 2). Of these 18 polar metabolites that changed in abundance in response to osmotic stress none were regulated by both SSU-1 and DAF-16 (Supplemental Table 2). In addition, we identified 63 lipid metabolites that exhibited a greater than 2-fold decrease in abundance in response to osmotic stress and 85 lipid metabolites that exhibited a greater than 2-fold increase in abundance in response to osmotic stress. Of these 148 lipid metabolites that change in abundance, six were regulated by both SSU-1 and DAF-16 and all six were lysophosphatidylcholines (Fig. 4c). Specifically, we identified six lysophosphatidylcholines that exhibited between two and eight fold increases in abundance in response to osmotic stress and this increase in abundance required both SSU-1 and DAF-16 (Fig. 4c). These findings show that embryonic metabolism is affected by osmotic stress and suggest that changes in lysophosphatidylcholine metabolism are involved in the response to osmotic stress.

In short, we have identified a neurohormonal signalling pathway that regulates insulin sensitivity and specificity in *C. elegans.* Specifically, we found that neurohormonal signalling via SSU-1 was required for the specificity of developmental arrest in response to osmotic stress, but not starvation, and that the loss of neurohormonal signalling via SSU-1 could suppress the enhanced sensitivity of insulin-like receptor (*daf-2*) mutants to osmotic stress. The finding that SSU-1 is required to control the expression of DAF-16 target genes in response to osmotic stress sheds light on how the inhibition of insulin-like signalling by both osmotic stress and starvation can cause different states of arrested development. We propose that SSU-1 functions in parallel to the FOXO transcription factor DAF-16 to promote a transcriptional response to osmotic stress, the production of lysophophatidylcholines, and developmental arrest in response to osmotic stress (Fig. 4d). We speculate that the regulation of development and modification of insulin signalling specificity by cytosolic sulfotransferases and nuclear hormone receptors is a conserved process and that studies of how cytosolic sulfotransferases affect metabolism and insulin specificity will provide important insights into disorders that involve insulin signalling, including obesity and type-2 diabetes.

## Materials and Methods

### Strains

*C. elegans* strains were cultured as described^8^ and maintained at 20°C unless noted otherwise. The Bristol strain N2 was the wild-type strain. Mutations used were:

**LGI:** *daf-16(m26)*, *daf-16(mu86)*

**LGIII:** *daf-2 (e1370)*

**LGIV:** *ssu-1(fc73)*, *ssu-1 (n5883)*, *ssu-1 (tm1117)*, *ssu-1(gk266317)*, *ssu-1 (gk747222)*, *ssu-1 (gk876992)*, *ssu-1 (gk319712)*, *zls356 [Pdaf-16::daf-16a/b-gfp; rol-6]*

**LGX:** *nhr-1 (n6217)*, *nhr-1(n6219)*, *nhr-1(n6228)*, *nhr-1(n6231)*, *nhr-1 (n6242)*

**Unknown linkage:** *wuls57 [sod-5p::GFP, rol-6(su1006)]*

**Extrachromosomal arrays:** *nEx2685 [Ptrx-1::ssu-1::mCherry::unc-54 3^'^UTR; Punc- 122::GFP], nEx2722 [Prab-3::nhr-1::tbb-2 3’UTR], nEx2720 [Pmyo-3::nhr-1::tbb-2 3’UTR], nEx2719[Pges-1::nhr-1::tbb-2 3’UTR]*

### M cell division in response to starvation

Analyses of M cell division were performed as described^2^.

### DAF-16::GFP localization

Embryos were placed onto Petri plates containing 500 mM NaCl in NGM agar seeded with *E. coli* OP50 for 5 hrs. Confocal microscopy was performed using a Zeiss LSM 800 instrument. The resulting images were prepared using ImageJ software (National Institutes of Health). Image acquisition settings were calibrated to minimize the number of saturated pixels and were kept constant throughout the experiment.

### Assay for developmental arrest

Approximately 200 developing eggs from mothers grown at 50 mM NaCl (unless otherwise noted) were collected and placed on standard NGM Petri plates containing varying concentrations of NaCl for 48 hrs. After 48 hrs, animals that remained immobile and were not feeding were scored as arrested. Mobile animals that were feeding were scored as developing. Percent failing to arrest was defined as the percent of animals mobile and feeding (unlike animals normally arrested in response to osmotic stress) and includes L1-stage larvae.

### Mutagenesis screen for animals that fail to arrest development

Approximately 20,000 L4 stage wild-type animals were incubated with 20 μl of ethyl methanesulfonate (Sigma) in 4 mL of M9 for 4 hrs at 20°C. F3 generation embryos were placed onto Petri plates containing 500 mM NaCl in NGM agar and screened for mutants that hatched and were mobile.

### Metabolite preparation and quantification

Approximately 100 μl of embryos were placed on standard NGM agar plates. After 3 hrs embryos were collected in M9, pelleted, and frozen. 100 μl of frozen embryos were resuspended in 600 μl methanol and lysed using a BeadBug microtube homogenizer (Sigma) and 0.5 mm Zirconium beads. After lysis, 300 μl of water and 400 μl of cholorform was added to each sample and samples were then vortexed for 1 min at 4°C and centrifuged for 10 min at 15000 g at 4°C. After centrifugation the polar and lipid layers were separated and dried using a SpeedVac concentrator. Liquid chromatography and mass spectrometry were performed as described^9^.

### RNA-Seq

Wild-type, *ssu-1 (fc73)*, and *daf-16(m26)* embryos were placed on standard NGM plates. After 3 hrs embryos were collected in M9, lysed using a BeadBug microtube homogenizer (Sigma) and 0.5 mm Zirconium beads (Sigma), and RNA was extracted using the RNeasy Mini kit (QIAGEN). RNA integrity and concentration were checked using a Fragment Analyzer (Advanced Analytical), and libraries were prepared using a Illumina NeoPrep RNAseq kit. Libraries were quantified using the Fragment Analyzer (Advanced Analytical) and qPCR before being loaded for paired-end sequencing using the Illumina NextSeq500. 75-nucleotide paired-end sequencing reads were mapped against the *C. elegans* genome assembly celO using RSEM v. 1.2.15, with Bowtie v. 1.0.1 for read alignment (flags --paired-end -p 6 --bowtie-chunkmbs 1024 --forward-prob 0, for strand-specific libraries)^10-11^. Expected read counts per gene were retrieved and, after rounding them to the nearest integer, used to perform differential gene expression with DESeq2 in the R v. 3.2.3 statistical environment^12-13^. Sequencing library size factors were estimated for each library; to account for differences in sequencing depth and complexity among libraries, as well as gene-specific count dispersion parameters (reflecting the relationship between the variance in a given gene’s counts and that gene’s mean expression across samples). Differences in gene expression between conditions (expressed as log2-transformed fold-changes in expression levels) were estimated under a general linear model (GLM) framework fitted on the read counts. In this model, read counts of each gene in each sample were modeled under a negative binomial distribution, based on the fitted mean of the counts and aforementioned dispersion parameters. Differential expression significance was assessed using a Wald test on the fitted count data (all these steps were performed using the DESeq() function in DESeq2)^13^. The DESeq2 algorithm also proposes two filtering steps to remove noisy outliers, one named “independent filtering”, based on the mean of normalized count, and a second one, based on the Cook’s distance, which assess how much the fit of the distribution would be impacted by removing an individual sample^13-14^. The results were the same in the presence or absence of these two filtering steps for the N2 vs. *ssu-1* comparisons. For the N2 vs. *daf-16* experiment, these two filtering steps were turned off for the purpose of identifying differentially-expressed genes, out of a concern that they would lead to the undue elimination of noisy, yet real differentially-expressed genes in that samples series. P-values were adjusted for multiple-comparison testing using the Benjamini-Hochberg procedure^14-15^ and genes with adjusted p-value < 0.05 and absolute fold-change greater than 2 were used for downstream analyses. To compare different mutants against wildtype, subsets of genes meeting the above-mentioned differential expression criteria in each mutant-wildtype comparison were intersected and enumerated Gene identifiers were retrieved from Wormbase version WS235. Hierarchical clustering and visualization of the data was performed in Spotfire (Tibco), using a complete linkage function and correlation as a distance metric both for genes and samples.

### Psod-5::GFP images

Embryos were placed onto Petri plates containing either 50 mM or 500 mM NaCl in NGM agar. Arrested L1 stage animals at 500 mM NaCl were imaged after 24 hrs (Fig. 2b and 2c). Early L1 stage animals at 50 mM NaCl were imaged after 5 hrs to control for staging (Fig. 2a and 2g). Dauer animals were collected 7 days after starvation on plates containing 50 mM NaCl (Fig. 2f). Confocal microscopy was performed using a Zeiss LSM 800 instrument for Figures 2b and 2c. Confocal microscopy was performed using a Leica SP8 instrument for Figures 2f and 2g. The resulting images were prepared using ImageJ software (National Institutes of Health). Image acquisition settings were calibrated to minimize the number of saturated pixels and were kept constant throughout each experiment.

### Cloning of Ptrx-1::SSU-1::mCherry

A synthetic *ssu-1* DNA fragment with synthetic introns replacing endogenous introns (endogenous introns are repetitive and could not be synthesized) was obtained from Integrated DNA Technologies using their custom gene synthesis service. The *ssu-1* fragment was amplified with a 3’ 5xGly linker using appropriate primers. The pJDM169 vector containing 1.1 kb of the *trx-1* promoter sequence upstream of the *trx-1* start codon, mCherry and an *unc-54* 3’UTR was obtained from J. Meisel (Meisel and Kim, 2014). The *ssu-1* DNA fragment with 5xGly linker was cloned into the pJDM169 using Infusion HD (Clontech) cloning to generate the plasmid pVDlOO that contains *Ptrx-1: :ssu-1(cDNA)* – *5xGly* – *mCherry::unc-54 3’UTR*.

### Cloning of nhr-1 rescuing transgenes

*R09G11.2c* was amplified from *C. elegans* wild-type cDNA and used in all vectors generated for tissue-specific expression of *nhr-1* cDNA. Infusion HD (Clontech) cloning was used to done individual promoter fragments with *nhr-1* cDNA, 5xGly linker, mCherry and *tbb-2 3’UTR* to generate plasmids containing the generic sequence *Promoter::nhr-1 cDNA* – *5xGly* – *mCherry::tbb-2 3’UTR*. The promoter fragments of the genes *ges-1* (in pVD104), *unc-54* (in pVD105), *rab-3* (in pVD106) and *dpy-7* (in pVD107) used for tissue-specific expression of *nhr-1* cDNA contain 2.9, 1.9,1.4 and 1.3 kb, respectively, of sequence upstream of each of these genes’ start codons.

### Statistics and Reproducibility

Two-tail t-tests were used to compare all samples that reflect percentages of populations or populations of animals. No statistical method was used to predetermine sample size. The experiments were not randomized. The investigators were not blinded to allocation during experiments and outcome assessment. Statistics source data available in Supplemental Table 4.

## Acknowledgments

We thank David Gems and the *Caenorhabditis* Genetic Center, which is funded by the NIH National Center for Research Resources (NCRR), for providing strains; N. An for strain management; and A. Doi, and A. Corrionero for helpful discussions. HRH, VD, KB, and NOB were supported by NIH grant GM024663 and NOB was supported by NSF grant 1122374. VD was a Howard Hughes Medical Institute International Student Research fellow. LRB and REWK were supported by NIH grant GM117408.

---

**Figure S1.**
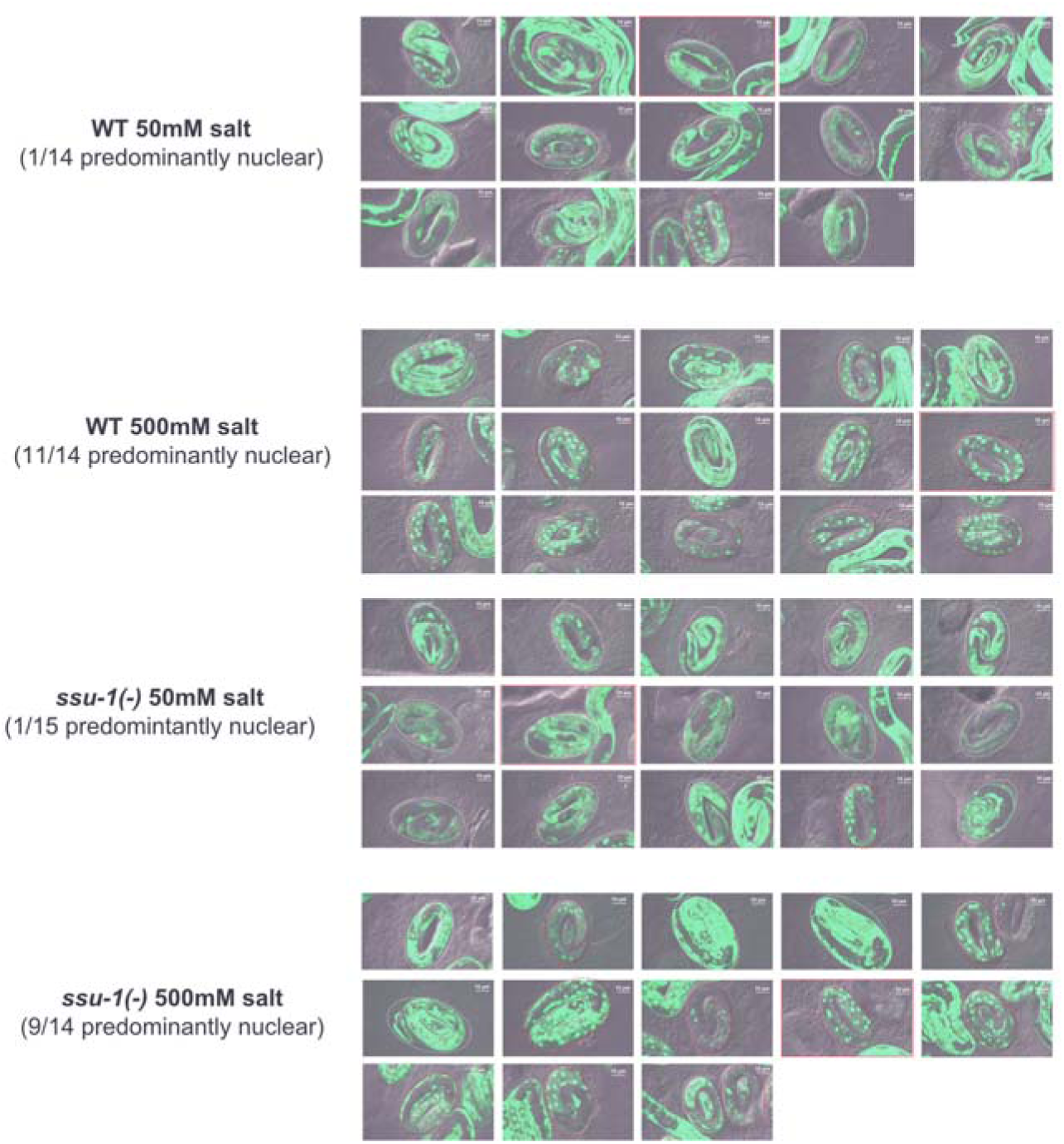
SSU-1 is not required for DAF-16 translocation into the nucleus in response to osmotic stress. Confocal images of DAF-16::GFP localization after 5 hrs of exposure to 500 mM NaCl in the wildtype and *ssu-1(fc73)* mutants. Scale bars, 10 μm. The red dashed lines indicate embryos scored as predominantly nuclear. The white dashed lines indicate embryos scored as predominantly cytoplasmic. Images with a red border were used as representative images in Fig. 3f.

*Supplemental Table 1* Profile of mRNA expression in wild-type and *ssu-1(fc73)* embryos at 50 mM NaCl and 500 mM NaCl.

*Supplemental Table 2* Profile of mRNA expression in wild-type and *daf-16(m26)* embryos at 50 mM NaCl and 500 mM NaCl.

*Supplemental Table 3* Profile of lipid and polar metabolites in wild-type, *ssu-1(fc73)*, and *daf-16(m26)* embryos from parents grown at 50 mM and 300 mM NaCl.

*Supplemental Table 4* Statistics source data.

